# A unifying framework for joint trait analysis under a non-infinitesimal model

**DOI:** 10.1101/293803

**Authors:** Ruth Johnson, Huwenbo Shi, Bogdan Pasaniuc, Sriram Sankararaman

## Abstract

**Motivation:** A large proportion of risk regions identified by genome-wide association studies (GWAS) are shared across multiple diseases and traits. Understanding whether this clustering is due to sharing of causal variants or chance colocalization can provide insights into shared etiology of complex traits and diseases.

**Results:** In this work, we propose a flexible, unifying framework to quantify the overlap between a pair of traits called UNITY (Unifying Non-Infinitesimal Trait analYsis). We formulate a Bayesian generative model that relates the overlap between pairs of traits to GWAS summary statistic data under a non-infinitesimal genetic architecture underlying each trait. We propose a Metropolis-Hastings sampler to compute the posterior density of the genetic overlap parameters in this model. We validate our method through comprehensive simulations and analyze summary statistics from height and BMI GWAS to show that it produces estimates consistent with the known genetic makeup of both traits.

**Availability:** The UNITY software is made freely available to the research community at: https://github.com/bogdanlab/UNITY

**Supplementary information:** Supplementary data are available at *Bioinformatics* online.

## 1 Introduction

Genome wide association studies (GWAS) have identified thousands of regions in the genome that contain variants that contribute to risk for many diseases. Many of these risk regions have been implicated in multiple phenotypes such as autism and schizophrenia (Autism Spectrum Disorders Working Group of The Psychiatric Genomics Consortium *et al*., 2017), multiple autoimmune diseases (Cotsapas *et al*., 2011; Ramos *et al*., 2011; Richard-Miceli and Criswell, 2012), Crohn’s disease and psoriasis (Ellinghaus *et al*., 2012), and many others. Understanding which causal variants are shared among diseases can provide novel etiological insight as well as provide evidence of potential shared causal mechanisms between complex traits. In addition, identifying which variants contribute to multiple traits can help decipher which molecular traits (e.g., gene expression) contribute to disease risk (Giambartolomei *et al*., 2014; Hormozdiari *et al*., 2016); genetic variants that causally alter gene expression as well as disease risk can link a particular gene to a given disease.

Genetic overlap has been analyzed both at the genome-wide level and local level, where the latter refers to analysis done within a given genomic region. *Genetic correlation*, a measure that quantifies the similarity in the genetic effects on pairs of traits, is commonly used for assessing the relationship between two traits and can be applied either genome-wide or to local data (Bulik-Sullivan *et al*., 2015; Shi *et al*., 2017). Many of the models for estimating genome-wide genetic correlation assume an *infinitesimal* genetic architecture where all SNPs are assumed to have a very small effect on the trait. In contrast to genetic correlation, *colocalization* methods aim to estimate whether the GWAS association signals for two traits at the same region are due to the same causal variant across the traits or chance(Giambartolomei *et al.,* 2014; Hormozdiari *et al.,* 2016). The methods that relax the infinitesimal assumption either assume a single causal variant per region or limit the number of potential causal variants a priori, often due to computational considerations (Giambartolomei *et al.,* 2014; Hormozdiari *et al.,* 2016). Although, both genetic correlation and colocalization aim to describe the genetic sharing between traits, these methods have been utilized largely independently of each other.

In this work we present a unifying statistical model that ties together genetic correlation and colocalization. To accomplish this, we present a fully generative Bayesian statistical model that models the shared as well distinct genetic variants underlying a pair of traits. The model allows for sparse genetic architectures (where only a small fraction of variants are causally impacting the traits). The model is richly parametrized: allowing us to jointly model global parameters such as the proportion of variants that are causal for both as well for either trait, the trait heritability, the correlation of the effect sizes at the causal SNPs, as well as local parameters such as the effect of a single SNP on each of the traits.

A challenge of a non-infinitesimal genetic architecture is that it presents a computationally challenging inference problem. Performing inference under this model often involves explicitly enumerating all causal configurations of the SNPs. This exponential search space of 2^2*M*^, where *M* is the number of SNPs analyzed, proves intractable given the large genetic data sets now available. We propose Unifying Non-Infinitesimal Trait analYsis (UNITY) that relies on Markov Chain Monte Carlo (MCMC) to approximate the posterior probabilities of the model parameters. In this work, we focus on estimating the proportion of shared and trait-specific causal variants since parameters such as heritability and genetic correlation can be estimated using previous methods (Bulik-Sullivan *et al.,* 2015). Additionally, a key advantage of the method is that it only requires summary level association statistic data, which bypasses many of the privacy concerns associated with individual level data. With the widespread availability of GWAS summary statistics (Pasaniuc and Price, 2017), we expect that a method operating only on summary statistics would prove most useful for the research community. Through comprehensive simulations and an analysis of height and BMI, we show that our method can accurately estimate the proportion of shared causal SNPs between two complex traits.

## 2 Methods

### 2.1 Generative Model

Here we introduce a Bayesian framework for estimating the proportion of causal variants shared between a pair of complex traits. The input to our method is the vector of signed effect sizes at each SNP for each trait (we only analyze SNPs for which effect size estimates are available for both traits). We model the genetic as well as non-genetic variances in each trait, the genetic correlation among the traits, and the proportion of causal SNPs that are shared across traits as well as are unique to each. The proportion of causal SNPs shared between the traits is denoted by *p*_11_, the proportion of causal SNPs specific to trait 1 and trait 2 as *p*_10_ and *p*_01_ respectively, and the proportion of non-causal SNPs is denoted by *p*_00_, where *p*_00_ + *p*_10_ + *p*_01_ + *p*_11_ = 1. For each trait *p* ∈ {1, 2}, we denote the genetic variance 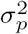 (which is the same as its heritability as 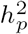 if the trait is standardized), the environmental noise as 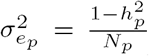, where *N*_*p*_ denotes the sample size for trait *p*, and the genetic correlation between the two traits as *ρ*. Altogether, our model has the following parameters: 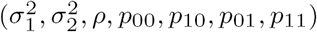.

We assume that trait *p* (*p* ∈ {1, 2}) measured in individual *i*, *y*_*p*,*i*_ is a linear function of standardized genotypes ***x***=(*x*_*i*,1_, …, *x_i_,M*) measured across *M* SNPs with SNP effect sizes *β*_*p*_ = (*β*_*p*_,1,…,*β*_*p*,*M*_) and independent additive noise term *∈*_*p*,*i*_. Further, we assume that there are no sample overlaps across the two studies.

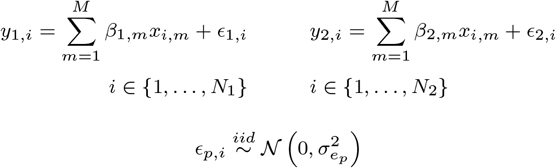

A SNP *m* is causal for trait p if its true effect *β*_*p*,*m*_ ≠ 0 and it is not causal otherwise. We denote the probability of a SNP being causal for every combination of the two traits as: *p* = (*p*_00_,*p*_10_,*p*_10_,*p*_11_).

Denoting the causal effect sizes for trait *p*, *p* ∈ {1,2} across all SNPs 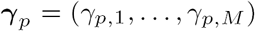, we assume that the causal effect sizes for each SNP are independent, allowing us to model the effect sizes at SNP *m* for each of the two traits 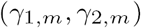 as a random vector drawn from a bivariate normal distribution centered at zero with the following covariance matrix:

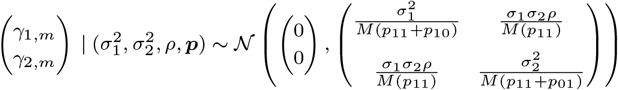

*C*_*p*_ = (*C*_*p*,1_,…, *C*_*p*,*M*_) denotes the causal indicator vector for trait *p*, where *C*_*p*,*M*_ = 1 if SNP *m* is causal for trait *p* and 0 otherwise. (*C*_1,*m*_, *C*_2_,_*m*_) is a random vector drawn from a discrete distribution with parameters given by *p*:

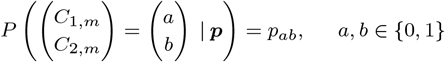

The true effect sizes for each trait *p* at SNP *m, β*_*p*,*m*_, conditioned on the causal status at a SNP is the element-wise product of the causal indicator vector and the true causal effect sizes.

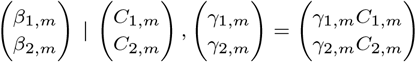

We can model the conditional distribution of the GWAS summary statistics given the true effect sizes, where 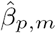 is the estimated marginal effect size of the *m*^*th*^ SNP for trait *p* (Shi *et al.,* 2017):

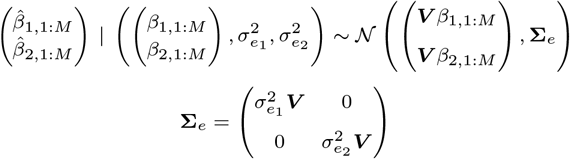

*V* is the matrix of correlations among the SNPs, *i.e.,* the linkage disequilibrium (LD) matrix. *V* can be estimated from a reference panel of genotypes collected from a population that is genetically similar to the populations for which summary statistics are available. Alternately, when performing inference at the genome-wide level, we can prune the list of SNPs such that they come from independent LD blocks. LD-pruning creates an approximately independent subset of SNPs in which case *V* can be approximated by the identity matrix, ***I***. In this work, we restrict our attention to the case where ***V*** ≈ ***I***.

We impose a Dirichlet prior on *p:*

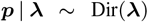

Here ***λ*** = (*λ*_1_, *λ*_2_, *λ*_3_, *λ*_4_). In practice, we set*λ*_1_=*λ*_2_ = *λ*_3_ = *λ*_4_ = *λ* = 0.20:

In principle, we can also impose priors on the remaining parameters, *i.e.,* the trait heritability 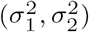 and their genetic correlation *ρ* and estimate all of these parameters jointly with *p* in a fully Bayesian model. These parameters can be estimated using other methods (Bulik-Sullivan *et al.,* 2015) and, in this work, we fix the values of these parameters to their estimates and focus on estimating *p.*

Given the parameters 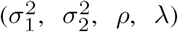, the joint distribution of the probability of causal configurations *p,* the causal indicator vectors *C*_1_, *C*_2_, the causal effect sizes 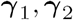 and the estimated effect sizes 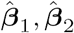 is given by:

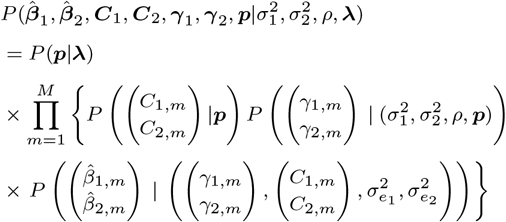

Integrating over the hidden variables *C*_1_; *C*_2_, *γ*_1_; *γ*_2_, we obtain:

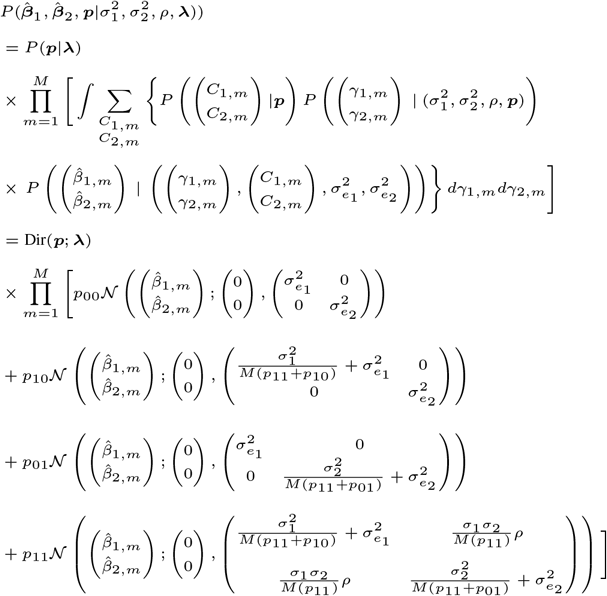

### 2.2 Parameter inference in our model

Given the generative model described in the previous section, the inference problem lies in computing the posterior distribution of *p* given the estimated summary statistics

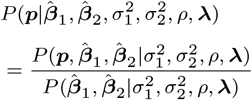

The true joint posterior distribution is intractable. Thus, we use Markov chain Monte Carlo (MCMC) (Brooks *et al.,* 2011) to approximate the posterior distribution. MCMC approximates the target posterior distribution

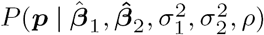 by a sequence of random samples 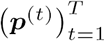 drawn from a Markov chain constructed so that the stationary distribution of the chain is the target posterior.

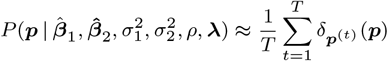

In our setting, we use a random-walk Metropolis-Hastings algorithm (Metropolis *et al.,* 1953) that generates a sample *p*^(*t*+1)^ at iteration *t* + 1 given the sample *p*^(*t*)^ at the previous iteration using the following proposal distribution that generates a proposed sample *p\** which is then accepted or rejected depending on the Metropolis-Hastings ratio (which depends on the ratio of the posterior probability density at the proposed parameter to the previous parameter):

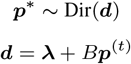

Here *B* is a constant that controls the variance of the proposal distribution. In practice, we found that *B =* 10 yields effective mixing.

The final step in specifying the MCMC algorithm lies in computing the ratio of the posterior probability density at the proposed parameter to the original parameter. Computation of the ratio requires the evaluation of the posterior probability only up to a normalization constant:

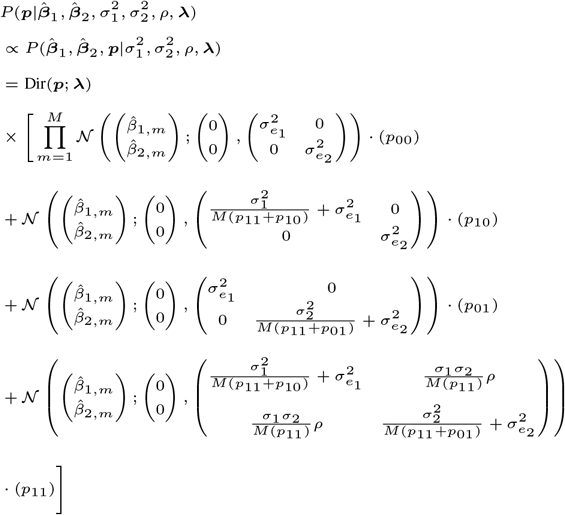

### 2.3 Efficient mixing of MCMC chains

In any practical application of MCMC, the number of iterations, burn-in period, and initialization point are critical to ensuring convergence and accurate estimates. Slow mixing of the MCMC chains can occur if the starting point is at a region of low posterior density. As opposed to selecting a random starting point, we carefully select the initialization of each chain by choosing the set of parameters that yields the highest posterior density. We use the Limited-memory BFGS algorithm (Byrd *et al.*, 1994) to determine the maximum a posteriori estimates for *p*_00_, *p*_10_, *p*_01_, *p*_11_. We repeat this 10 times, initializing the optimization algorithm with random starting values drawn from the prior. We compute the posterior density of all 10 candidate starting values and select the set that yields the highest density. This set of parameters is then used as the starting point for our MCMC chain. In addition, to diagnose convergence, we use 100 Markov chains all initialized using the scheme described above. Our final estimate is the mean of all samples drawn from the 100 chains.

**Table 1.** Displayed is a summary of current methods that perform joint trait analysis and the relationship to the parameters in UNITY. Boxes with an (*) denote the values that a method models. Note that this summary is not exhaustive

| Method | $h^2$ | $\rho$ | $p$ | misc. |
| --- | --- | --- | --- | --- |
| UNITY | * | * | * |  |
| Cross-trait LD Score regression (Bulik-Sullivan *et al.*, 2015) | * | * | $p_{11} \approx 1$ | |
| PleioPred (Hu *et al.*, 2017a) | * | * | * | infers $p$ to estimate effect sizes |
| COLOC (Giambartolomei *et al.*, 2014) | — | — | * | max 1 causal |
| eCAVIAR (Hormozdiari *et al.*, 2016) | — | — | * | max 6 causals |

### 2.4 Note on runtime

We assessed the performance based on the number of seconds per iteration of the MCMC sampler. The main computation is calculating the likelihood at each iteration, which is directly dependent on the number of SNPs per trait. The complexity of the algorithm is 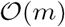, where *m* is the number of SNPs. We empirically demonstrate that our method is linear in the number of SNPs through simulation (Supplementary Figure 1). In addition, the runtime is invariably connected to the number of iterations required for the MCMC to converge. We find that using the maximum a posteriori probability (MAP) estimate as an initialization value leads to fast convergence, requiring only 500 iterations in practice.

## 3 Results

UNITY provides a novel generalized framework to jointly model GWAS summary statistics data of two complex traits, incorporating fundamental genetic parameters, such as heritability and genetic correlation, and makes minimal assumptions in inference procedures. Since UNITY assumes a non-infinitesimal model, it allows for very sparse genetic architectures, i.e. by setting *p*_00_ ≈ 1. However, this non-infinitesimal model can also be generalized to the infinitesimal model by setting *p*_00_ ≈ 0,*p*_10_ ≈ 0, *p*_01_ ≈ 0,*p*_11_ ≈ 1.

### UNITY generalizes colocalization and genetic correlation

We discuss a comparison of the parameters of UNITY with those obtained by other methods that perform cross-trait analysis and the underlying assumptions of each method. We first analyze the cross-trait LD score regression model (Bulik-Sullivan et al., 2015), which estimates genome-wide genetic correlation based on the random-effect model, making the implicit assumption that every SNP has a non-zero effect. In contrast to cross-trait LD score regression, UNITY assumes a generalized non-infinitesimal model, explicitly modeling a sparse genetic architecture. We also compare UNITY with methods that do not make the infinitesimal model assumption. While models such as PleioPred explicitly model the proportion of trait-specific and shared causal variants *p*_00_, *p*_10_, *p*_10_, *p*_11_, the main goal of this method is to perform genetic risk prediction (Hu *et al*., 2017a) rather than estimating these proportions.

We compare UNITY with COLOC (Giambartolomei et al., 2014) and eCAVIAR (Hormozdiari *et al*., 2016), Bayesian methods to assess the evidence of colocalization, i.e. whether GWAS signals of two traits are driven the same underlying causal variants. Both methods explicitly model *p* = (*p*_00_, *p*_10_, *p*_10_, *p*_11_) (Giambartolomei *et al*., 2014; Hormozdiari *et al*., 2016). However, COLOC makes the simplifying assumption that thereisat most one causal variantat a region (Giambartolomei *et al*., 2014), allowing it to not explicitly model LD. And although eCAVIAR allows for multiple causal variants and explicitly models LD,itrestricts the maximum number of causal variants at 6 per region for computational efficiency (Hormozdiari *et al.,* 2016). In comparison with these methods, UNITY allows for any number of causal variants while making the assumption that there is no LD between the SNPs. We outline a summary of the relationship between UNITY and all methods described in Table 1.

To empirically demonstrate the benefit of the relaxed assumptions of UNITY as compared to current methods, we conduct a modest comparison against COLOC (Giambartolomei *et al.,* 2014). We simulated 100 regions of 500 SNPs with multiple causal variants. We perform colocalization analysis over all of the regions using COLOC. When there are causal variants independently associated with each trait and shared variants, COLOC estimates that the association within the region is driven only by two independent variants, where one is specific to trait 1 and the other is specific to trait 2. Because COLOC assumes at most one causal variant per region, the method is unable to distinguish between a variant that independently drives only one trait versus a variant that is colocalized when both cases are present. For completeness, we also included a simulation that follows the assumption underlying COLOC of the one-causal setting. The full table listing these results in outlined in Supplementary Table 2. However, we are unable to directly compare estimates with COLOC because there is not a clear mapping between the estimates of COLOC and the estimated parameters of UNITY, thus any direct comparison would be an unfair comparison due to the mismatch in the models.

### 3.1 Simulations

We generated summary statistics for 500 SNPs from two synthetic GWAS. The causal effect sizes for each SNP, 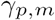, were drawn jointly from a multivariate normal distribution where 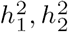, *ρ* denote the heritability of each trait and the genetic correlation. We denote the number of SNPs as *M* and the proportion of causal variants for each trait as *p*_10_, *p*_01_ and the proportion of shared casuals as *p*_11_:

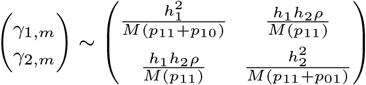

To simulate causal SNPs, we drew an *M* × 4 matrix from a multinomial distribution parametrized by *p* where the *m*^*th*^ row of values denotes whether a SNP is causal for neither trait, only trait 1, only trait 2, or neither trait. Using this, we constructed two *M* × 1 causal indicator vectors, *C*_1_, *C*_2_, where *G*_1,*m*_, *G*_2,*m*_, = 1 if the *m*^*th*^ SNP was causal for both traits, C_1,m_ = 1, *C*_2,*m*_ = 0 if the SNP was only causal for trait 1, *C*_1,*m*_ = 0, *C*_2,*m*_ = 1 if it was only causal for trait 2, and *C*_1,*m*_, *C*_2,*m*_ = 0 if the SNP was non-causal. To get the true effect sizes, we multiplied element-wise *β*_1_ = *C*_1_ o γ_1_ and *β*_2_ = *C*_2_ o γ_2_ where we are essentially zeroing out any entry from the causal effect vector where a SNP is non-causal.

To compute the estimated GWAS effect sizes, 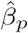, we assumed cov(*ϵ*_1_, *ϵ*_2_) = 0, so random noise terms *ϵ*_1_, *ϵ*_2_ were drawn from two normal distributions 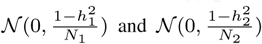 respectively. We assume that the SNPs being used at the genome-wide level will be LD-pruned such that there is very little or no correlation structure. Thus, we set the LD matrix *V* = *I*_*M*_. where *I*_*M*_ is an M × M identify matrix. We then draw the estimated effect sizes from a conditional distribution of the GWAS summary statistics, as described in Methods.

First, we confirm that our method accurately predicts the proportion of causal variants under varying sample sizes and heritability estimates. We tested a variety of simulation frameworks where we fixed the genetic correlation and heritabilities of the two traits. We ran each simulation for 500 iterations and used the first quarter of the iterations as burn-in. We vary the proportion of causal variants contributing to only trait 1 (*p*_10_), proportion of causal variants for only trait 2 (*p*_01_), and the proportion of casual variants contributing to both traits (*p*_11_). As shown in Figure 1., we can see that UNITY performs robustly across each scenario.

**Fig. 1.**
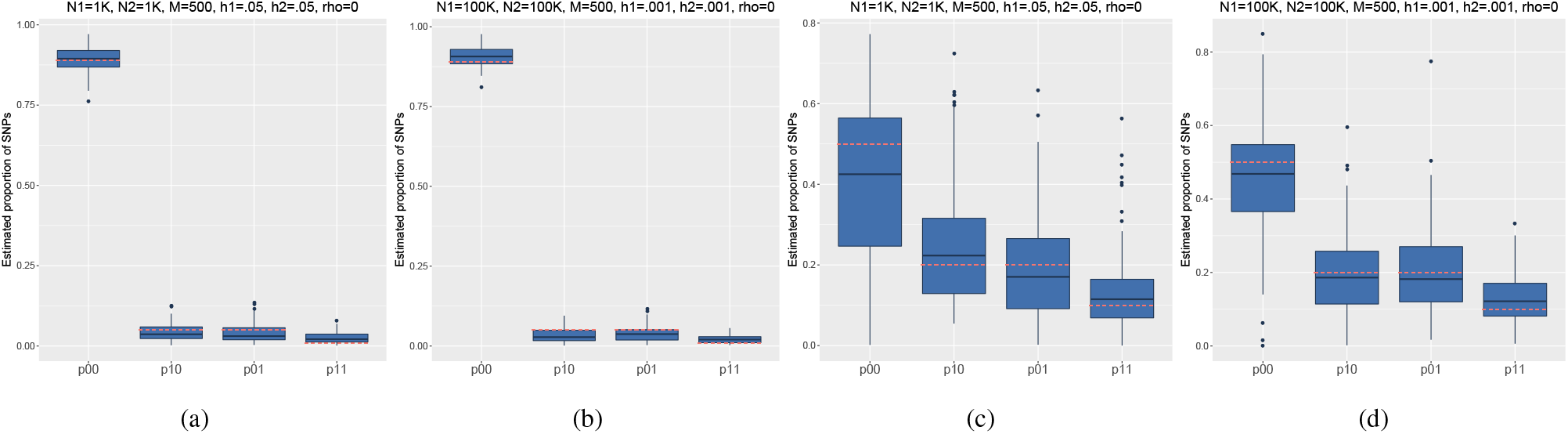
We estimate the proportion of causal variants under four simulation frameworks where we vary the sample size (*N*_1_, *N*_2_), heritability 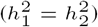, and proportion of causal variants. First, we first simulated values where the total proportion of causal variants is low: *p*_00_ = 0.89, *p*_10_ = 0.05, *p*_01_ = 0.05, *p*_11_ = 0.01, along with a low sample size and high heritability: 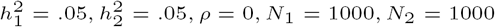, as shown in (a). Second, we tested the model with the same proportion of causal variants, but with a larger sample size and smaller heritability: 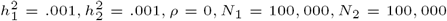, shown in (b). Third, we simulated data with a higher proportion of causal variants, *p*_00_ = 0.50, *p*_10_ = 0.20, *p*_01_ = 0.20, *p*_11_ = 0.10. Using the same sets of heritabilities and sample sizes from the first two simulations, we tested the prediction accuracy of our model. (c) denotes the simulation with low sample size and high heritability, and (d) denotes the simulation with high sample size and low heritability. The dotted red lines denote the true proportion of causal SNPs in each simulation.

Next, to assess how UNITY performs with varying levels of heritability, we continued to fix *ρ* = 0, but varied the values of the heritability. Note that we used low heritability values due to the low number of simulated SNPs (M=500). From Figure 2, we can see that the estimates reflect the prior distribution of (*p*_00_, *p*_10_, *p*_01_, *p*_11_) when the heritability is very low. We also show in Figure 3 that our estimates are invariant to the correlation between phenotypes.

**Fig. 2.**
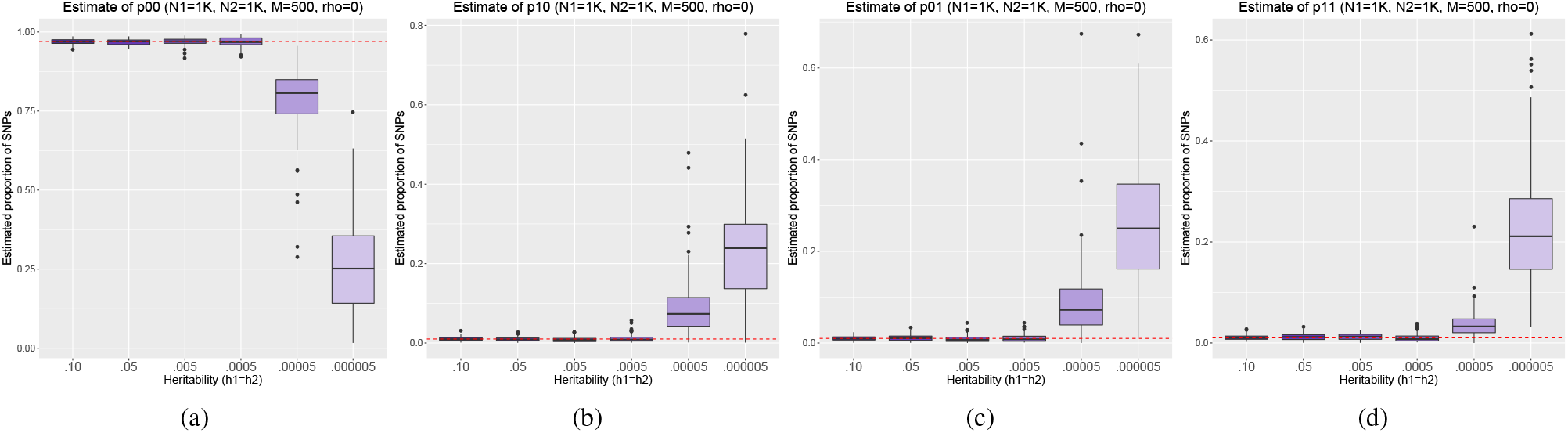
We simulate the following proportion of causal variants *p*_00_ = 0.97, *p*_10_ = 0.01, *p*_01_ = 0.01, *p*_11_ = 0.01 and vary the heritability 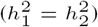 while fixing *ρ, N*_1_, *N*_2_, *M.* We vary the heritability from .01 to 5e-7 and plot the estimated proportion of non-causal variants (a), proportion of causal variants for trait 1 (b), proportion of causal variants for trait 2 (c), and proportion of shared causal variants (d). We note that as the heritability goes down, the data becomes less informative and the estimates reflect the prior.

To assess the role of sample size in our inference, we performed simulations where we varied the number of individuals from 1,000 to 250,000. We find that the recommended sample size should be at least 50,000 individuals to yield precise results (Supplementary Figure 2). Additionally, to further assess the performance of the method, we also performed simulations where 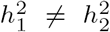 and when 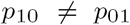. Through simulation, we demonstrate that our method is robust to these scenarios, with detailed results provided in Supplementary Figure 3 and Supplementary Figure 4.

Finally, through simulations, we empirically demonstrate that our method is well calibrated under the null hypothesis, defined as: (1) *p*_10_ = 0, (2)*p*_01_ = 0, and(3)*p*_11_ = 0. To demonstrate this, we simulated 100,000 SNPs with 100,000 individuals where 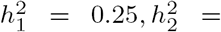 0.25, *ρ* = 0. For each hypothesis, we set the parameter of interest exactly to 0 and then then simulated 2% causal variants between the remaining parameters. For example, for null hypothesis (1), the corresponding set of simulation parameters would be: *p*_10_ = 0, *p*_01_ = 0.01, *p*_11_ = 0.01. Using UNITY, we estimated the null parameter and report the posterior mean and standard deviation in Table 2. Note that UNITY estimates the null parameter very close to zero, but not exactly zero. This is because there is a nonzero prior on the set of parameters, making it not possible to be exactly zero, but can instead be asymptotically close.

**Table 2.** We present the posterior means and standard deviations estimated when the proportion of causal variants is set exactly to zero for trait 1 and trait 2, and when the shared proportion is exactly zero.

| Hypothesis | Null parameter | Mean | SD |
| --- | --- | --- | --- |
| 1 | $p_{10}$ | 0.0006 | 0.0023 |
| 2 | $p_{01}$ | 0.0004 | 0.0005 |
| 3 | $p_{11}$ | 0.0002 | 0.0003 |

### 3.2 LD pruning to identify approximately independent SNPs

To rigorously assess the role of LD in our model, we demonstrate a sufficient LD-pruning scheme through simulations. To model arealistic LD structure, we used SNPs from 1000 Genomes (Consortium *et al.,* 2012a) to compute the LD for each of the approximately independent LD blocks identified in Berisa et al (Berisa and Pickrell, 2016). We filtered rare SNPs with MAF ≤ 0.05 and used 1 million SNPs sampled across the LD blocks. We chose only a subset of 1 million SNPs because this closely reflects the number of SNPs genotyped on SNP arrays. We simulated the GWAS effect sizes as outlined in Section 3.1, where the heritabilities for each of the each traits was set to 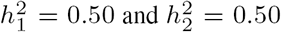 (which is similar to the estimated SNP heritability for height), and genetic correlation *ρ* = 0.

**Fig. 3.**
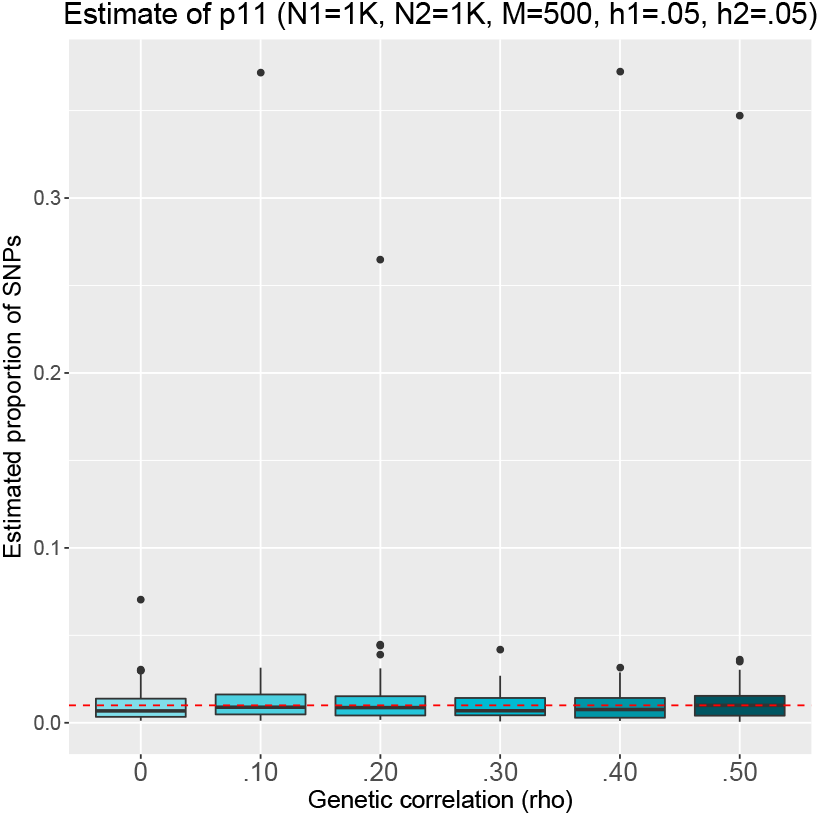
We simulate the following proportion of causal variants *p*_00_ = 0.97, *p*_10_ = 0.01, *p*_01_ = 0.01, *p*_11_ = 0.01 and vary the genetic correlation from 0 to 0.50 while fixing 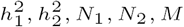. We only show the estimate of *p*_11_ since this would be the only estimate directly affected by the presence of genetic correlation.

**Fig. 4.**
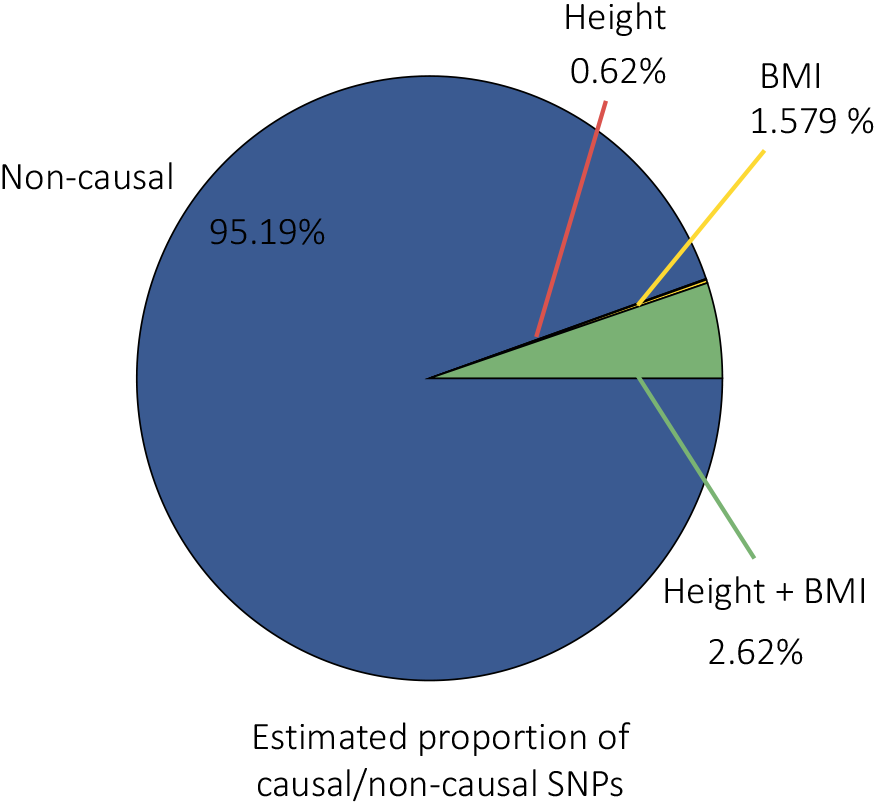
We show the distribution of estimated non-causal and causal SNPs from the Height and BMI analysis.

To assess the role of LD-pruning, we divided the genome into K kilobase non-overlapping windows and selected a SNP from each window. We varied K to assess the minimal window size necessary to create a subset of approximately independent SNPs. In addition, we used cross-trait LD Score regression to estimate the heritabilities for both traits and the genetic correlation after pruning, which were subsequently used in the inference. Through simulations, we determined that a 5KB window provides precise estimates (Supplementary Table S1).

### 3.3 Empirical analysis of BMI and Height

We downloaded GWAS summary data for both Height and BMI from the GIANT consortium (Speliotes *et al.,* 2010; Allen *et al.,* 2010) where each study has > 170, 000 individuals. First, we overlapped each GWAS by rsid to get SNPs present in both studies. Then for each trait, we filtered out SNPs with a minor allele frequency ≤ 0.05. Additionally, we performed LD-pruning by taking a SNP from every 5KB window.

We used cross-trait LD Score to estimate the heritability and genetic correlation parameters: 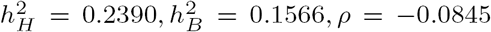. Denoting Height as the first trait and BMI as the second, we estimated the proportion of causal variants for each trait as, *p*_00_ = 0.9519, *p*_10_ = 0.0062, *p*_01_ = 0.01579, *p*_11_ = 0.0262. We summarize the distribution of estimated causal SNPs in Figure 4.

Our results are consistent with the known genetic makeup of BMI and Height. Since BMI is a function of an individual’s height and weight, we expect all of the contributing variants for Height to also contribute to BMI. UNITY predicts more BMI-only specific variants than Height-only variants. We hypothesize that the BMI specific variants are those that contribute to weight, whereas the variants that contribute to height in the BMI dataset were already captured in the *p*_11_ estimate. In principal, we would expect *p*_10_ to be zero since SNPs contributing to height also contribute to BMI. We expect this could be due to the nonzero prior on *p*_10_. Because of this, the estimate can never truly be zero but can be asymptotically close.

## 4 Discussion

In this work, we introduce a statistical framework for quantifying the relationship between two complex traits. The key advantage of our method is that it makes very few assumptions about the data and few restrictions during inference. Rather than relying on assumptions about a trait’s genetic architecture, we let the data describe the underlying genetics. By using a Metropolis-Hastings sampling framework, we can calculate a variety of likelihoods without relying on any conjugate prior pairings. For example, although we choose to model the causal effect sizes through a multivariate normal, one could choose another distribution, and the sampling procedure would still hold even if the new distribution did not have a conjugate prior. Finally, by operating exclusively on GWAS summary statistic data, we aim to encourage future large-scale meta analyses, since obtaining individual level data is not always readily available.

We conclude with several limitations and potential future directions of our framework. First, as the size of genetic datasets grow, subsampling methods such as MCMC may prove computationally intractable. Alternatives include using adaptive MCMC to accelerate mixing and convergence or variational methods that do not require subsampling. Additionally, we have yet to rigorously quantify the effects of LD in our model in practice for local inference. We leave rigorous comparison between UNITY and other relevant methods as future work.

Additionally, recent integrative methods have shown that the incorporation of a variants functional genomic context can improve both power and accuracy in identifying potential causal variants (Pickrell, 2014; Kichaev *et al*., 2014; Li and Kellis, 2016; Hu *et al*., 2017b). Large-scale initiatives such as the ENCODE (Consortium *et al*., 2012b) and ROADMAP (Roadmap Epigenomics Consortium *et al*., 2015) projects have provided comprehensive databases of tissue-specific functional genomic annotations. Combining this rich atlas of functional data and the genetic information from GWAS will likely uncover novel insights into disease biology. We leave the incorporation of functional elements as a potential direction for future work.

## Acknowledgments

We are grateful to Kathryn Burch, Claudia Giambartolomei, and Gleb Kichaev for helpful and insightful discussions.

